# CactEcoDB: Trait, spatial, environmental, phylogenetic and diversification data for the cactus family

**DOI:** 10.1101/2025.06.27.661607

**Authors:** Jamie B. Thompson, Catherine Martinez, Jorge Avaria-Llautureo, Santiago Ramírez-Barahona, Gerardo Manzanarez-Villasana, Alastair Culham, Andrew Gdaniec, George Ryan, Chris Venditti, Georgia Keeling, Nicholas K. Priest

## Abstract

Integrated datasets linking traits, spatial distributions, environmental variables and phylogenies are essential for comparative research, but remain limited for many plant taxa, including those which are most threatened. Cactaceae are a morphologically and ecologically diverse succulent family that are iconic components of ecosystems across the Americas, and face high extinction risk. To support future comparative research, we present CactEcoDB (The Cactus Ecological Database), an Open Access dataset of curated spatial, ecological, trait, phylogenetic, and diversification data for over 1,000 cactus species. CactEcoDB includes species-level trait data, geographic occurrence records, environmental variables, range size estimates, speciation rates, and the largest time-calibrated phylogeny of cacti to date. By integrating these diverse data in a single platform, CactEcoDB enables analyses of trait evolution, trait–environment interactions, biogeography, climatic niche evolution, and macroevolutionary dynamics in one of the most celebrated and threatened plant families.

## Background & Summary

Recent research highlights the importance of comparative methods for our understanding of the origins and ecological drivers of global biodiversity in large families of plants ^1–12^. This type of comparative research relies on comprehensive datasets that adequately capture the complexity of trait evolution, environmental variation, and lineage diversification ^13–35^. There is a growing appreciation of how this complexity, especially ecological interactions, coevolutionary dynamics, and environmental heterogeneity, shapes macroevolutionary and macroecological trajectories ^9,10,12,36–41^.

As global change accelerates and data availability expands unevenly across lineages, so does the need for integrative, high-quality datasets for threatened and unique groups^25,26,42–50^. However, assembling such datasets remains difficult and time-consuming, often requiring expert curation and synthesis across disparate data types and sources.

The cactus family (Cactaceae) is among the most morphologically and ecologically distinctive plant lineages ^51–57^. Comprising approximately 1,850 species, cacti are found in an extraordinary range of habitats, from hyper-arid deserts and tropical dry forests to high-elevation grasslands and volcanic islands ^58^. Centers of species richness are found in Mexico, the Andes, and Eastern Brazil ^51–57^, but their distribution extends from southern Canada to Patagonia, with one species probably occurring naturally outside of the Americas (*Rhipsalis baccifera*) ^59^. Cacti are well known for their highly specialised morphological and physiological adaptations to water-limited environments, including stem succulence, heavily modified leaves, thick cuticles, and crassulacean acid metabolism (CAM) photosynthesis ^10,12,51,54^. These traits make them a classic example of adaptive evolution in plants, and in particular, a model system for understanding evolutionary responses to extreme environmental conditions.

Comparative research in the Cactaceae has bearing on diverse fields including plant physiology ^60–62^, biogeography ^51,63–66^, ecology ^67–69^, evolutionary dynamics ^10,12,51,54^, and conservation biology ^53,66,70–73^. The growing clarity of phylogenetic relationships within the family ^52^ has facilitated tests of the evolution of key traits such as growth forms ^60^, pollinators ^51,56^, and chromosome count ^74^. The power to detect drivers of diversification is greatly aided by the remarkably wide distribution and range of growth forms found in diverse environments across gradients including aridity, temperature, and elevation ^51,52,55,74^.

Beyond their ecomorphological adaptations, cacti hold exceptional cultural and economic significance ^75–77^. Archaeological evidence shows that species such as *Opuntia ficus-indica* were consumed more than 9,000 years ago in Mesoamerica ^78,79^, where they continue to serve as important sources of food, forage, and dye ^80–82^. Today, cacti are cultivated globally for ornamental purposes, and their remarkably varied forms, ranging from tiny globose species to massive columnars ^60^, make them among the most collected and traded groups in horticulture ^83^ along with orchids. They are unusual in horticulture in that the bulk of trade is of natural species, not highly bred and modified cultivars. The deep cultural history, combined with their horticultural desirability, has led to intense pressure on wild populations through unsustainable collection and illegal trade ^84,85^. As a result, Cactaceae is one of the most threatened plant families. A 2015 assessment found that 31% of cactus species are threatened with extinction, with habitat loss, climate change, and overexploitation cited as primary drivers ^70^. More recent projections suggest that 60-90% may experience future range contractions under near-future climate change scenarios ^71^.

The integration of phylogenetic, trait, and spatial data at the species level in cacti has remained fragmented, limiting the ability of researchers to address broad ecological and evolutionary questions, or to support conservation assessments grounded in comprehensive ecological data ^72,86^. To fill this gap, we present CactEcoDB, a curated dataset that integrates spatial, ecological, trait, and phylogenetic information for over 1,000 species of Cactaceae. This is a greatly updated and expanded version of previous datasets ^10^. CactEcoDB includes biotic trait data (e.g., growth form, plant height and pollinator), expert-curated geographic ranges, numerous important environmental variables, and an updated phylogeny, the largest time-calibrated molecular phylogeny for the group. As compared with the largest previous dataset ^10^, CactEcoDB expands and improves the available data: i) it increases in species sampling for plant height by 24.93%, spatial variables by ∼6%, and adds eight new variables; ii) it expands the number of classified growth forms from two to nine; iii) it replaces highly incomplete GBIF data with expert ranges defined by the IUCN ^53,66,70–73,87^, which were manually verified against independent data, especially the Plants of the World Online (POWO) ^88^; and iv) it provides a general-purpose resource for ecological, evolutionary, biogeographic, and conservation-focussed research. Furthermore, we have substantially expanded and updated several of the biotic traits with higher resolution and species coverage than previously available ^10^. By bringing together large scale data covering all aspects of cactus diversity in an Open Access resource, CactEcoDB enables new opportunities for research on trait-environment relationships, climatic niche evolution, and conservation, in one of the most unique and threatened plant families. Given the gradual increase of trait, spatial and phylogenetic data generally, we welcome the submission of additional data, which will improve future research into cacti.

## Methods

The methods presented here include a brief description of methods used to generate previous data ^10^, and our new efforts designed to improve and expand sampling of spatial distributions, environmental data, and traits (Figure 2). Although (10) is a large foundation of this work, we added ten new variables, recollected or added data to 39 traits (76.40% of CactEcoDB), and undertook thorough checking steps.

### Phylogenetic reconstruction

We provide two phylogenetic hypotheses. One is the original phylogeny estimated by Thompson et al. ^10^ (henceforth referred to as V1), and one is re-estimated using a phylogenomic constraint that has been published since ^89^. For both trees, the supermatrix approach was used, to maximise taxonomic sampling using publicly-available data in Genbank. 18 published plastid and nuclear loci were compiled from GenBank, by identifying and clustering orthologous sequences with the OneTwoTree pipeline ^90^, and aligned them using MAFFT ^91^ with quality inspection undertaken in SeaView ^92^. Partial sequences were merged with full-length sequences for each species where homologous sequences clustered into a fragmented and full cluster, retaining the longest version. Outgroup sequences from Anacampserotaceae, Portulacaceae, and Talinaceae were added using MAFFT’s “--add” function ^93^. After alignment, poorly aligned regions were trimmed using trimAl with the “gappyout” setting ^94^, and concatenated all loci into a supermatrix using AMAS ^95^. A Maximum Likelihood (ML) phylogeny was reconstructed with RAxML v8 ^95,96^, applying a GTR+G model partitioned by locus and assessing support with 300 bootstrap replicates. Time calibration was performed under Penalized Likelihood criteria with treePL, using secondary constraints on crown and stem age for the family derived from highest posterior densities from a relaxed-clock fossil-calibrated phylogeny of angiosperms ^5^ (see ^10^ for full details), in the absence of an informative fossil record of Cactaceae ^55^. Cross-validation was used to optimize the smoothing parameter across 100 priming runs, and the final dating was based on the smoothing value with the lowest score.

To make V2, we replicated these steps but constrained relationships in the Maximum Likelihood tree search according to a recently published phylogenomic backbone ^89^. We assessed relationships in this phylogenomic tree in the context of node support, and identified 15 nodes to constrain. These were tribes Blossfeldioideae, Cacteae, Cereeae, Copiapoeae, Cylindropuntieae, Fraileeae, Lymanbensonieae, Notocacteae, Opuntieae, Phyllocacteae, Pterocacteae, Rhipsalideae, and subfamilies Maihuenioideae, Pereskioideae and Leuenbergerioideae. This approach greatly improved relationships compared to the old phylogeny, given the low sequence variation found in cacti which commonly leads to weakly-supported nodes ^10,52,55^. The number of nodes supported by 70% bootstrap support in V2 (BS) increased from V1 by 11.8% (32.3 to 36.1%), and the number of nodes supported by 90% increased by 72.9% (11.9 to 20.6%). Eleven species displayed pathologically long terminal branches when the constraint was applied and were pruned.

### Diversification rate estimation

To estimate speciation rates from the phylogenies we used three methods: 1). Bayesian Analysis of Macroevolutionary Mixtures (BAMM) ^97^ (Figure 1), 2). Missing State Speciation and Extinction (MiSSE) ^98^, and 3). the DR statistic ^35^. Each of these have different assumptions and approaches, and associated benefits and pitfalls. BAMM is a Bayesian approach which discretises rates in the phylogeny into lineage-specific shift regimes, and provides a posterior sample to account for uncertainty. It also allows lineage-specific sampling fractions, which are of great importance given the phylogenetic imbalance of molecular sequencing across cactus groups ^10,52^. Although BAMM has received criticism, notably regarding sensitivity to priors and the accuracy of deep rate shifts ^99,100^, these have been defended statistically ^101,102^ and it remains one of the most accurate methods ^103^. MiSSE also discretises rates, but in a Maximum Likelihood framework providing only one estimate per model, and accounts for incomplete sampling with a global sampling fraction. While this does not account for within-group variation in sequencing efforts, lineage-specific sampling fractions have been criticised ^104^. Finally, the DR statistic allows rapid estimation as it only makes use of variation in terminal branch lengths and root-to-tip node count. However, it cannot account for incomplete and imbalanced sampling, and is known to be noisy compared to other metrics ^105^. We recommend using BAMM estimates, as BAMM accounts for incomplete sampling in a manner that does not assume that each lineage has been sampled to an identical fraction, as well as providing a posterior sample of estimates which accounts for uncertainty.

**Figure 1:**
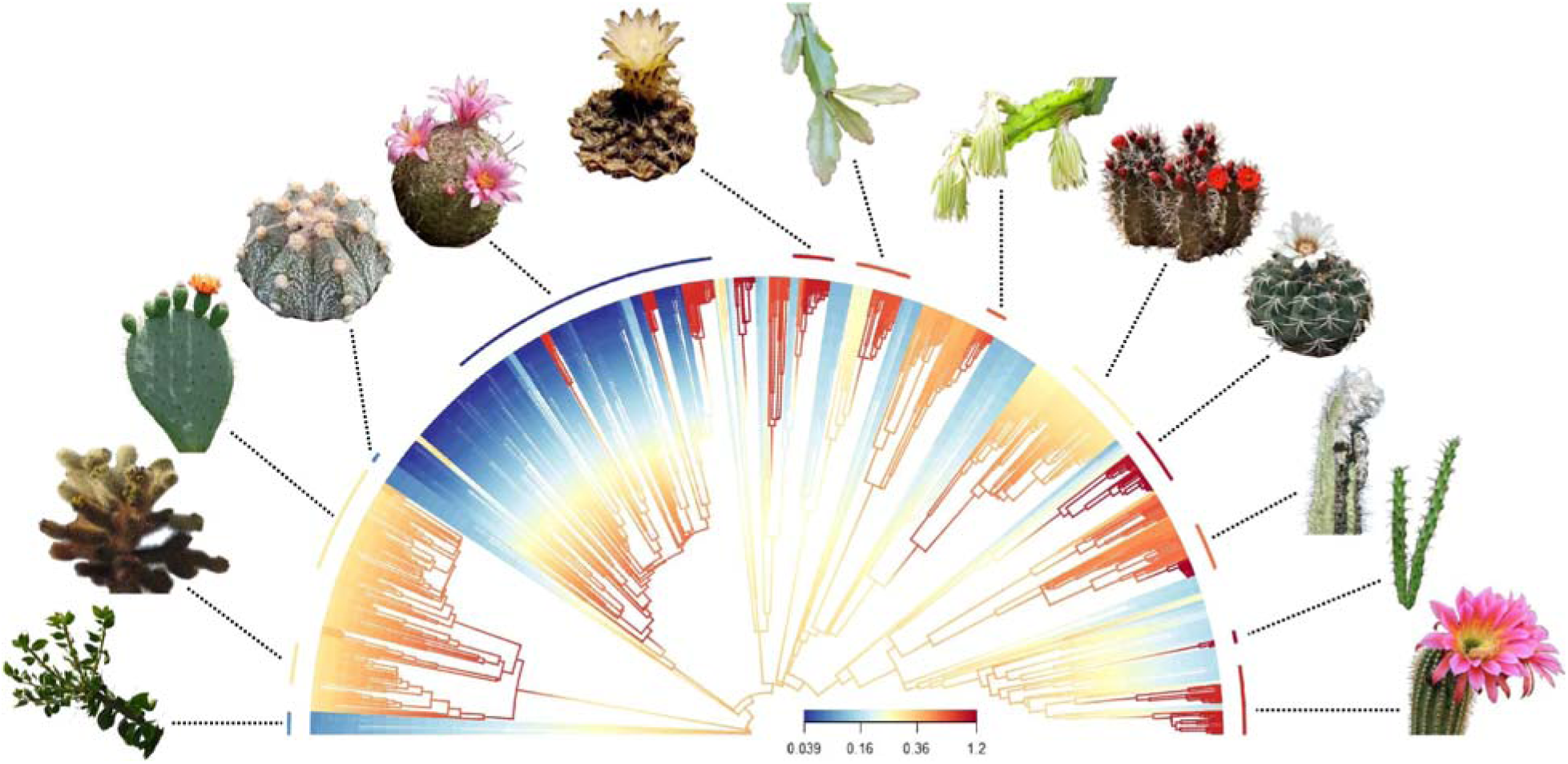
The phylogeny (V1), showing estimated speciation rate variation in one method (BAMM). Phylogenetic branches in phylogeny V1 ^10^ are coloured according to speciation rates estimated with BAMM ^97^, and vary 32-fold. Arc segments of median speciation rate for thirteen morphologically varied cactus genera are indicated. This figure and figure legend is reproduced with permission from ^10^. Cactus images are used under Creative Commons with modifications allowed. From left to right: images 1, 3, 8, 11, 12, and 13 used photos taken by Amante Darmanin, Forest & Kim Starr, John Tann, Renee Grayson, and Wendy Cutler, which are licensed under a Creative Commons Attribution 2.0 License (https://creativecommons.org/licenses/by/2.0/). Image 2 used a photo marked as being in the Public Domain (https://creativecommons.org/publicdomain/mark/1.0/). Images 4 and 10 used photos taken by Leonora Enking and Lyubo Gadzhev, which are licensed under a Creative Commons Attribution-ShareAlike 2.0 License (https://creativecommons.org/licenses/by-sa/2.0/). Images 5, 7 and 9 used photos marked as being in the Public Domain using the CC0 1.0 Universal Public Domain Dedication (https://creativecommons.org/publicdomain/zero/1.0/). Image 6 used a photo taken by Christer Johansson, which is licensed under a Creative Commons Attribution 3.0 Unported License (https://creativecommons.org/licenses/by/3.0/deed.en).

**Figure 2:**
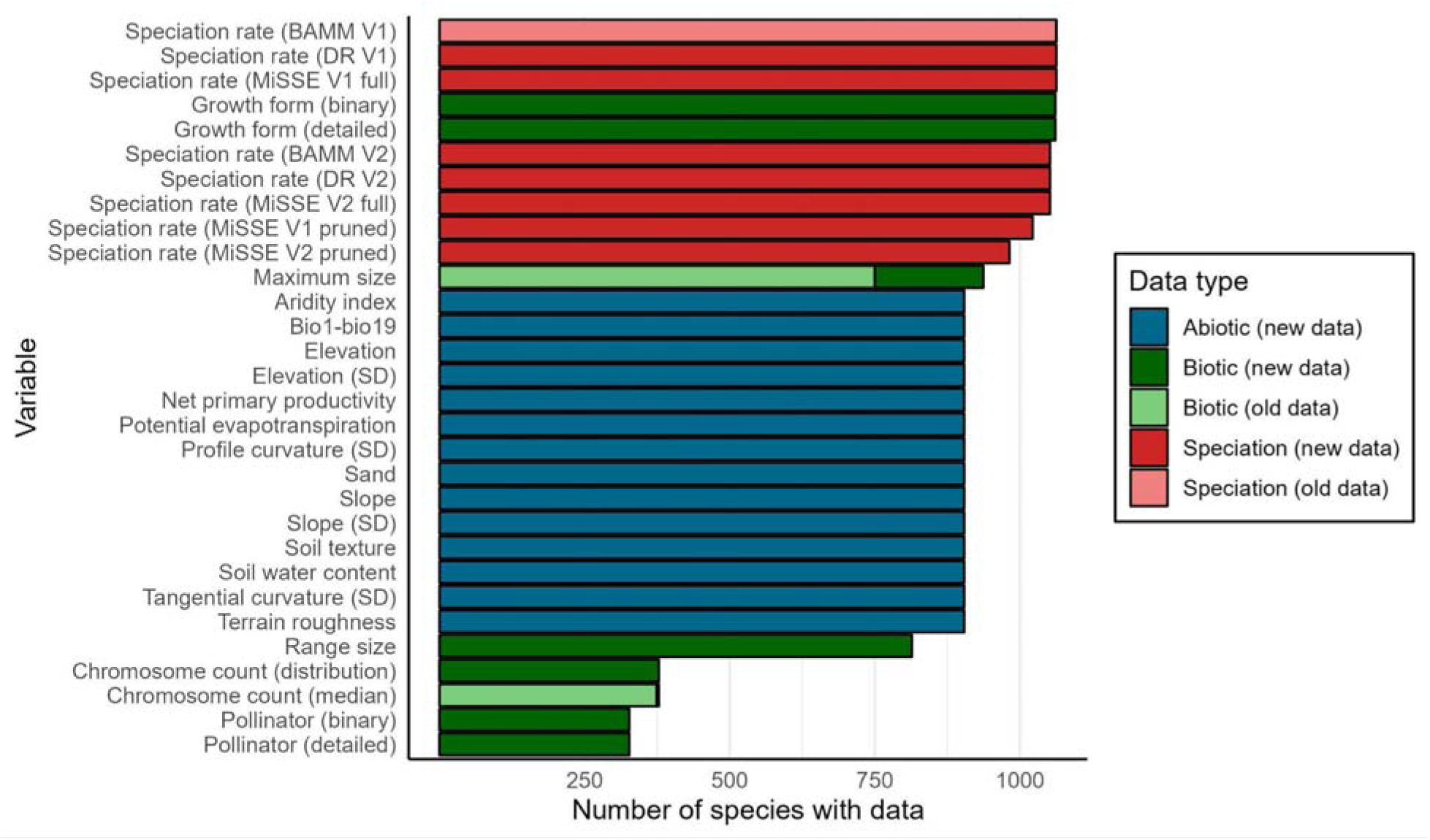
The distribution of species coverage by variable. Speciation rate estimates, growth form and plant size (maximum height or length) have highest coverage (1,063-982, 1,061 and 937 species, respectively), and the spatial environmental variables sample 904 species. We only calculated range size for those species in IUCN, given the occurrence scarcity in GBIF for the remaining species. Pollinator and chromosome count have lower coverage with 327 and 378 species, respectively. We compare the species coverage for variables in CactEcoDB against those in the largest previous dataset ^10^.

For phylogeny V1, we used the BAMM results from Thompson et al. ^10^. We replicated this process with V2. Prior to analysis with BAMM, outgroups were pruned. Incomplete sampling was accounted for in BAMM by providing genus-level sampling fractions. Four Markov Chain Monte Carlo (MCMC) chains were run for 50 million generations with BAMM, sampling every 5,000 generations and discarding the first 10% as burn-in.

Priors were set using the BAMMtools package in R ^106^, and a conservative prior allowing for a single rate shift was chosen to minimise overfitting. Convergence was assessed using the coda R package, ensuring effective sample sizes (ESS) of >200 for all key parameters. Following this, posterior distributions were summarized with BAMMtools, the best shift configuration was estimated for visualisation, and mean speciation rates at the tips were extracted for each species. As discussed in ^10^), we provide the entire event data file from BAMM to enable analysis of lineage diversification dynamics, with the caveat that exact divergence timings in cacti are highly uncertain due to the absence of informative fossils ^55^.

For the MiSSE analyses we used a global sampling fraction of 0.575 (phylogeny V1) and 0.572 (phylogeny V2), respectively, and estimated 30 possible model structures using the R package HiSSE ^107^, which were generated with the function generateMiSSEGreedyCombinations from combinations of between one and ten turnover parameters, and one and three extinction parameters. Using the function MiSSEGreedy, we estimated MiSSE parameters on each of the 30 models, then estimated ancestral diversification rates using the command MarginReconMisse.

Finally, we estimated tip rates using the function GetModelAveRates on the marginal reconstruction results. We replicated this on a version of each phylogeny that has the shortest branches pruned, using a threshold of 0.01 Mya, because formal-SSE models can be biassed by tiny branch lengths ^107,108^. In cacti, these could be real (recent divergences) or an artefact, and it is difficult to unambiguously classify them. In V1, 41 species were pruned and in V2, 70 species were pruned. We present results from both the unpruned and pruned phylogenies. For the DR statistic analysis, we used the R function DR_statistic (<u>github.com/Cactusolo/rosid_NCOMMS-19-37964-T</u>).

### Trait data compilation

We compiled species-level trait data from a combination of authoritative textbooks ^109^, primary taxonomic literature, specimen descriptions in online floras, and prior datasets ^10,51,60^. These data vary in their coverage across cactus species (Figure 2). Key traits that we updated, expanded, and standardised in our new dataset (i.e. beyond ^10^), include plant height, growth form, pollination syndrome, and chromosome count. We standardised traits previously recorded with different units, e.g. for height (to cm).

Where minimum and maximum plant heights were reported, we recorded maximum values to reduce biases from juvenile, diseased, or partial specimens, as is standard practice in comparative research into plant height ^110^. We expanded and updated all records for plant maximum size to improve species sampling by 187 species, from 750 to 937.

We provide data on the binary growth form scoring system developed by Hernández-Hernández et al. ^51^ and extended in sampling by ^10^. This system included two main categories grouping six subcategories: compact including smaller forms such as globose and barrel) and arborescent (containing larger forms such as columnar and shrubby).

However, we have devised a novel and more refined system using nine growth form categories for CactEcoDB: arborescent, barrel, columnar, cushion forming, epiphytic, geophytic, globose caespitose, globose solitary, and shrubby. For context, a previous growth form system used in (Hernández-Hernández et al., 2011) scored species as arborescent, shrubby, columnar, globose solitary, globose caespitose and barrel.

Briefly, arborescent cacti are those with a main trunk that branches above the base, typically with a tree-like structure; shrubby cacti lack a dominant trunk and instead exhibit basitonic branching from the base; columnar species are unbranched with elongated stems, including tall or prostrate cylindrical forms; globose solitary and globose caespitose refer to spherical-stemmed species under 0.5 m tall, growing singly or in clumps, respectively; barrel cacti have spherical stems exceeding 0.5 m in height; cushion-forming species grow in dense mats without a clearly assignable alternative form; geophytic species grow at or below ground level but are not cushion-forming; and epiphytic species are those growing on other plant hosts. We hypothesise that this system of nine growth forms adequately explains growth form variation in cacti, but note there is uncertainty and other systems exist ^60,71^. We scored these species based on botanical descriptions primarily drawn from ^109^, but also from other sources where unavailable or inadequately described in ^109^ (Figure 3).

**Figure 3:**
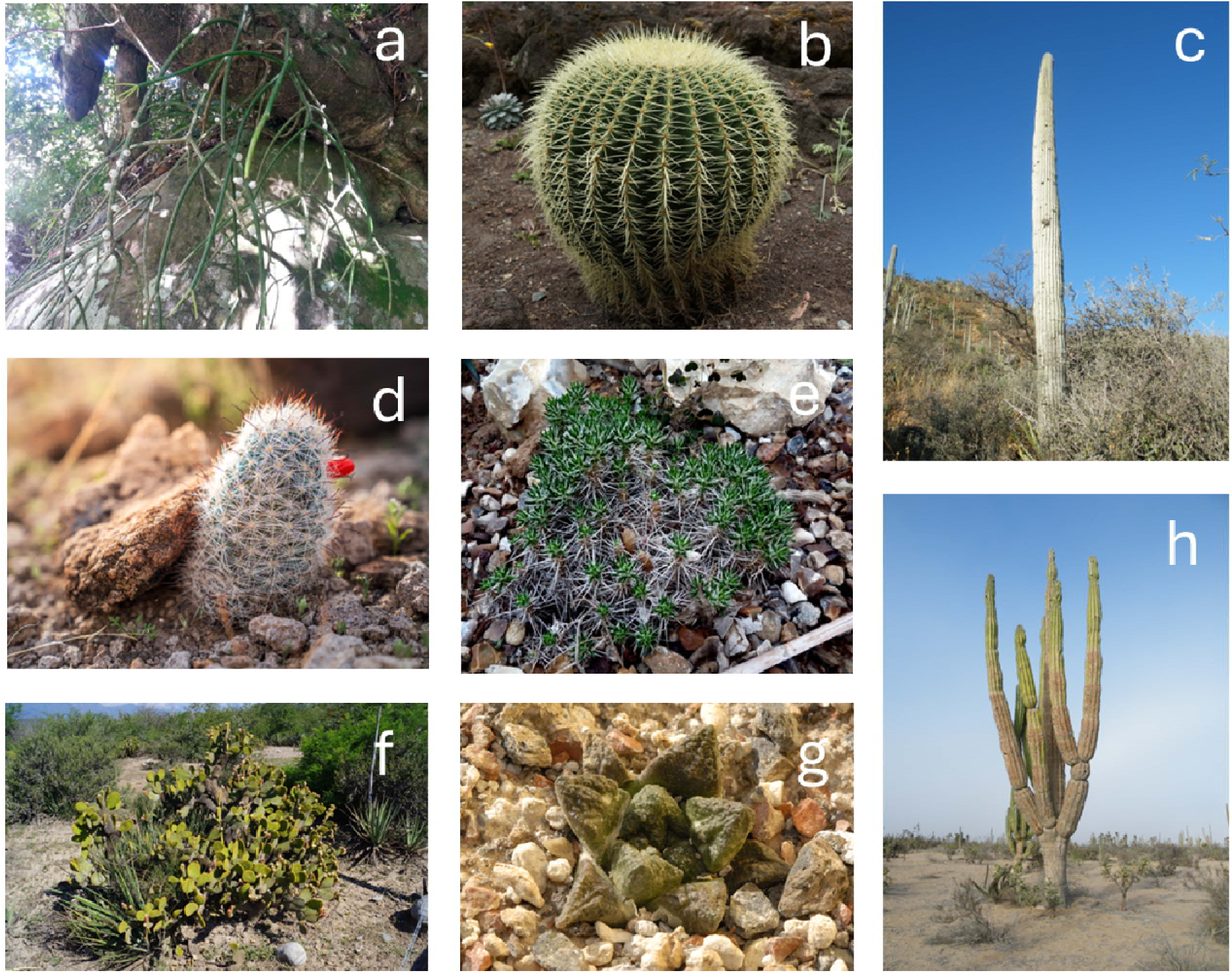
Examples of each growth form in CactEcoDB. (a) Rhipsalis baccifera, an epiphytic cactus. (b) Echinocactus grusonii, a barrel cactus. (c) Cephalocereus columna-trajani, a columnar cactus. (d) Mammillaria tetrancistra, a globose solitary species. Note that we have not shown a globose caespitose species, which appears as clusters of globose solitary species. (e) Maihuenia poeppigii, a cushion-forming species. (f) Opuntia microdasys, a shrubby species. (g) Ariocarpus fissuratus, a geophyte. (h) Pachycereus pringlei, an arborescent species. Cactus images are used under Creative Commons licenses with modifications allowed. Image (a) used a photo taken by Maria Vorontsova, which is licensed under the Creative Commons CC0 1.0 Universal Public Domain Dedication (https://creativecommons.org/publicdomain/zero/1.0/). Images (b) and (g) used photos taken by Dr. Hans-Günter Wagner, which are licensed under a Creative Commons Attribution-ShareAlike 2.0 License (https://creativecommons.org/licenses/by-sa/2.0/). Images (c) and (h) used photos taken by Amante Darmanin, which are licensed under a Creative Commons Attribution 2.0 License (https://creativecommons.org/licenses/by/2.0/). Image (d) used a photo taken by Jesse Pluim (BLM), which is marked as being in the Public Domain using the Public Domain Mark 1.0 (https://creativecommons.org/publicdomain/mark/1.0/). Image (e) used a photo taken by Laurent Houmeau, and image (f) used a photo taken by Sergio Niebla; both are licensed under a Creative Commons Attribution-ShareAlike 2.0 License (https://creativecommons.org/licenses/by-sa/2.0/).

We compiled pollination data and classified cactus species into five functional pollinator groups: bats, bees, birds (which are primarily Trochilidae), moths (which are primarily Sphingidae), and other insects (Figure 4). This level of resolution reflects the standard used in pollination research for cacti and other plant groups ^51,111–114^, and greatly improves on the binarisation approach (ancestral versus derived) established by ^10,51^. In botanical data, observations and inferences of pollinators are typically reported at taxonomic levels higher than species, as we have provided here, and lower taxonomic levels are inconsistent and rarer. The pollination syndromes themselves are shaped by shared floral adaptations that map reliably to higher-level clades (e.g., orders or functional groups) rather than to individual pollinator species ^115–117^. Species-level identification is uncommon in the botanical literature due to both data limitations and the diffuse, coevolutionary nature of many plant–pollinator interactions ^118,119^. To assemble these classifications, we combined information from existing compilations at genus, tribe and whole-family levels (e.g. ^120–122)^ with targeted literature searches using query combinations such as “genus + pollinator’’ and “genus + pollination.’’ All pollination assignments in ^10^, which used data primarily provided by ^51^, were re-assessed and 56 species were removed at the higher taxonomic level because their source could not be verified beyond ^51^. By systematically validating records against traceable literature and retaining only those with adequate support, our dataset provides a foundation for comparative research as well as future data collection. Our pollination data covers 17.68% of cacti, a larger proportion than a recent compilation for Orchidaceae (∼10%)51,114.

**Figure 4:**
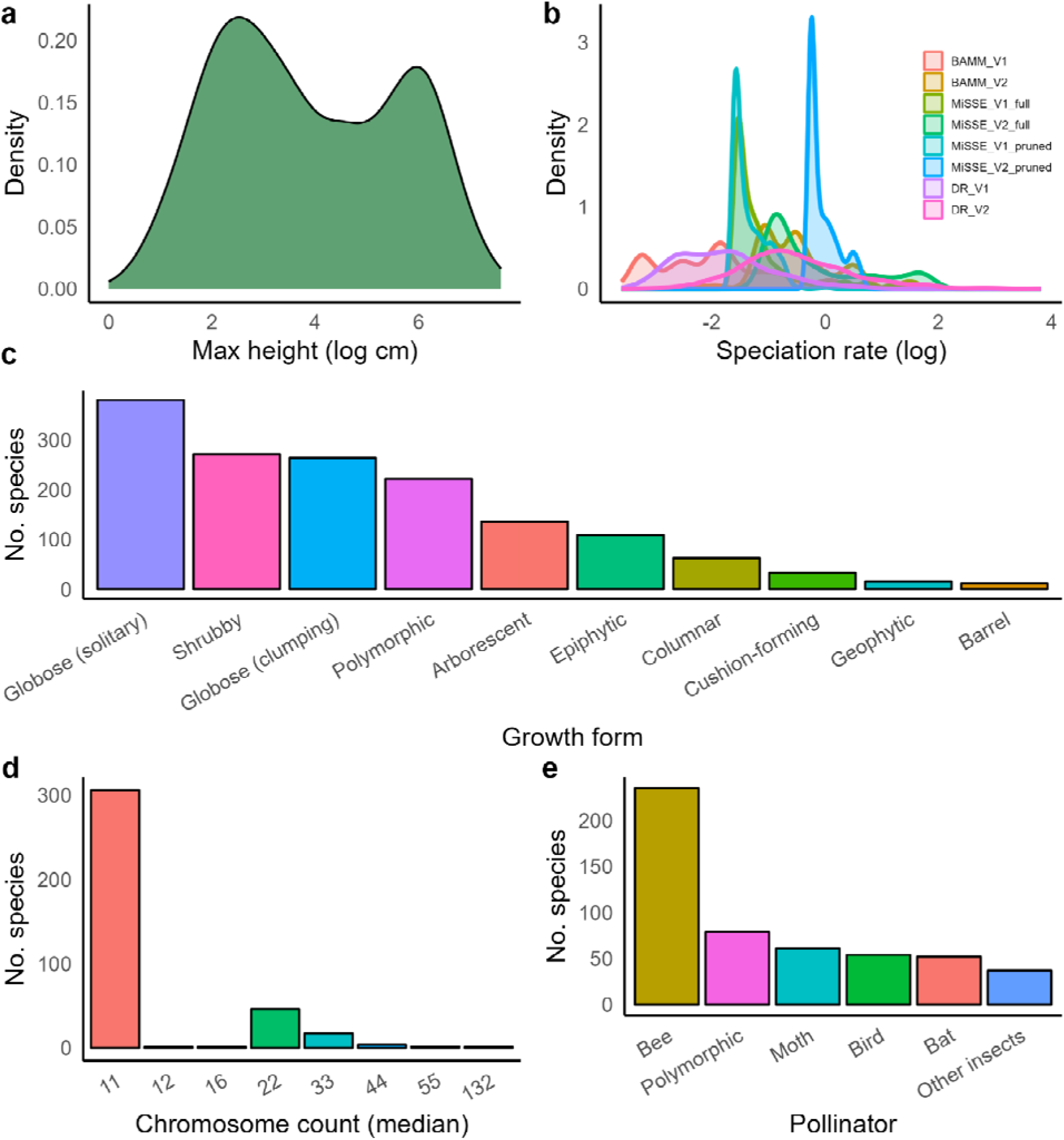
The distribution of plant height, speciation rate, growth form, chromosome count, and pollinator. (a) Cactus size (height or length) is bimodal, which is shaped by underlying growth form variation ^10^. (b) Tip speciation rate is mostly relatively slow, with fewer relatively faster species. (c) The most common growth form is globose solitary, followed by globose caespitose and shrubby. The least common growth forms are cushion forming, geophytic and barrel. 222 species are polymorphic, which mostly consists of species which are both globose solitary and globose caespitose (154 species), or both arborescent and shrubby (50 species). (d) Most species have the base number of 11 chromosomes, and most chromosome count variation is generated by polyploidy ^74^, reflected in the distribution (e.g. 22, 33, 44, 55). (e) Most species are pollinated by derived pollinators (bats, birds, moths) rather than the ancestral pollinators (bees) ^51^.

Finally, we captured data on chromosome counts from the Chromosome Count Database (https://ccdb.tau.ac.il/) ^123^, which compiled data from other sources and involved a cleaning step. We retained all recorded counts instead of relying on the reported medians (Figure 5). As well as allowing within-species variation to be captured, this allowed the correction of some zero median counts reported for species with chromosome counts in the database. However, we also replicate the median chromosome count for each species as it is useful for macroevolutionary research ^124–126^. We performed statistical and manual checks of collected trait data which flagged multiple records as spurious, which we either removed, confirmed or edited based on deeper literature searches (see Technical Validation).

**Figure 5:**
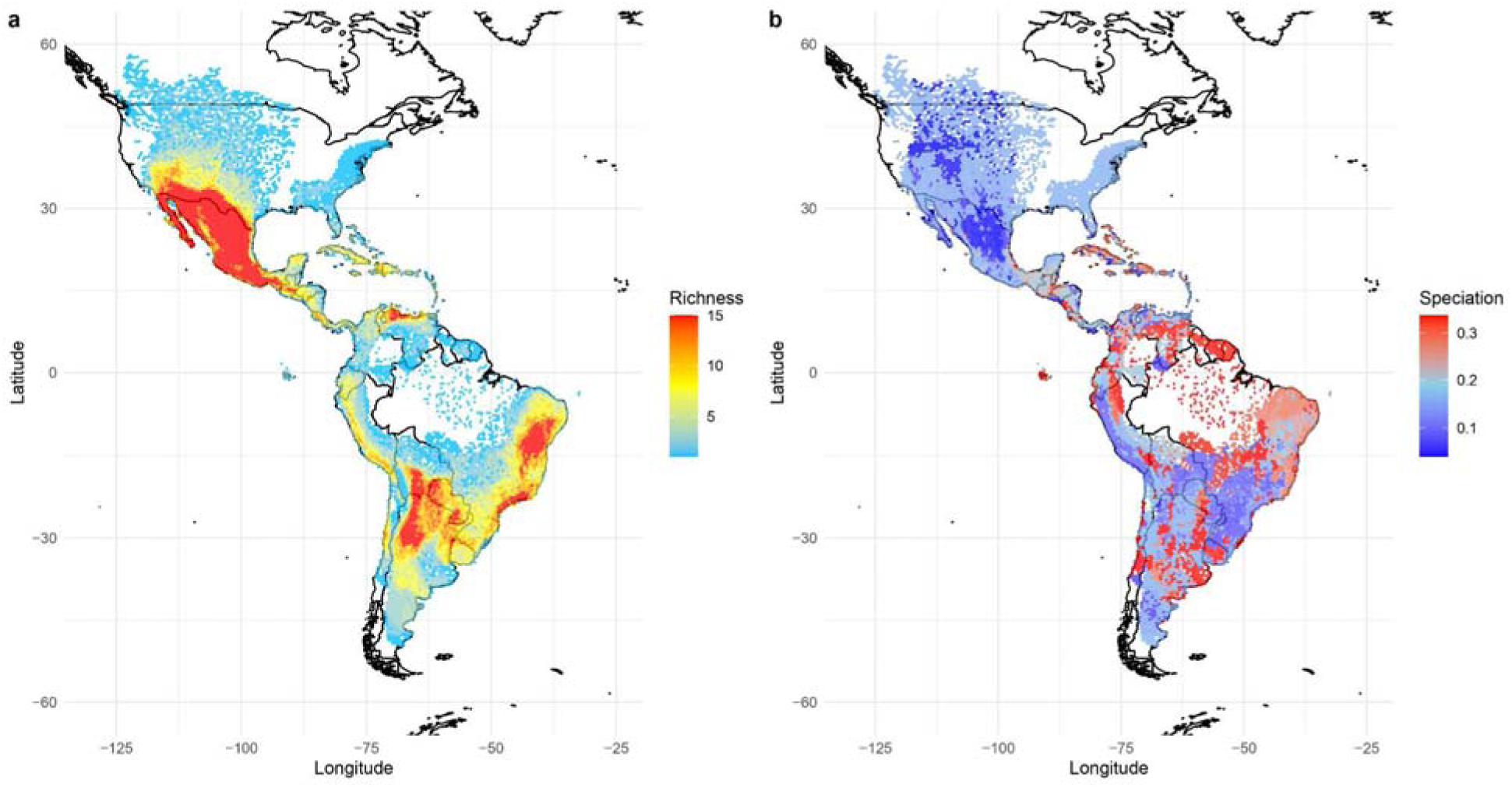
The spatial distribution of cactus richness and tip speciation rates. (a) Species richness is plotted per grid cell (∼50*50 miles), and shows Mexican, Eastern Brazilian and Andean richness hotspots. (b) Median tip speciation rate (BAMM_V1 from ^10^) is plotted per grid cell, showing a relative decoupling between areas of highest species richness and high speciation rate. This figure was presented in the supplementary materials of ^10^, but with lower resolution and different underlying spatial data.

### Distribution data

Previous distribution range datasets used data downloaded and cleaned from the Global Biodiversity Information Facility (GBIF, https://www.gbif.org/). However, we have greatly improved species and distributional sampling with a new sampling strategy (Figure 5). For CactEcoDB, we have primarily used expert range polygons available in the IUCN Red List of Threatened Species (https://www.iucnredlist.org/) ^70,127^, with some occurrences newly-acquired and curated from GBIF to maximise species sampling, following ^128^. This approach increases species sampling by 54 species (∼6%) and provides more accurate estimates of geographic range sizes, which allows us to fully characterise the environments of the entire ranges of species. Contrary to previous datasets, our new data reduce geographic bias, especially for species with poor GBIF records.

We downloaded IUCN ranges and manually checked them against the independent data sources available in POWO (https://powo.science.kew.org/) which records native and introduced ranges at a broad level ^88^. This identified one IUCN range that had sampled likely-introduced South American populations (for *Cylindropuntia tunicata*). We corrected this accordingly and then we sampled coordinates randomly from within each range with the number of coordinates sampled in relationship to the range size, following an established protocol ^128^. Specifically, we used a binning function to assign sample sizes based on range area (in km²): species with ranges <10 km² were sampled 20 times; 10–100 km², 50 times; 100–1,000 km², 150 times; 1,000–10,000 km², 300 times; 10,000–100,000 km², 500 times; 100,000–1,000,000 km², 700 times; 1,000,000–10,000,000 km², 900 times; and >10,000,000 km², 1,200 times. This sampling approach ensured adequate environmental representation for both narrow and wide-ranging species, while avoiding excessive data volume or downstream computational demands, particularly for the most widespread taxa.

We curated the most up-to-date GBIF occurrences, for those species not covered by the IUCN expert ranges. We used a rigorous and multi-step cleaning pipeline developed by ^129,130^. We queried all records under “Cactaceae”, considering only “Preserved Specimens” and removed species present in the IUCN expert range data. The code checked taxonomy against POWO and flagged suspect records for removal based on multiple criteria: i) records with missing or badly formatted coordinates; ii) records associated with biodiversity institutions; iii) records with equal or zero latitude and longitude; iv) records within a 10km radius of a country’s capital; v) records within 5km of a country’s political centroid; vi) records in the sea ^131^. As a final step, occurrence records that were outside the range described by POWO for each species were removed ^88^.

### Environmental data and verifying against GBIF data

We extracted spatial variables hypothesised to be important for cacti, succulents, and angiosperms generally (summarised in ^10^), from three different sources. From CHELSA ^132^ (https://www.chelsa-climate.org/), we extracted the 19 bioclimatic variables, aridity index, potential evapotranspiration and net primary productivity. From EarthEnv ^133^ (https://www.earthenv.org/), we extracted elevation and six measures of topographic complexity including slope, roughness, standard deviation of slope, and profile curvature. From SoilGrids ^134^ (https://isric.org/), we extracted sand content, water availability, and texture. We calculated median values per species for continuous variables, since they are less likely to be influenced by outliers, and also provide the entire distribution for each species across their range, to capture intraspecific variation. The latter is especially valuable when analysing trait–environment relationships, because it accounts for environmental heterogeneity within species ranges, instead of relying on a point estimate. Incorporating intraspecific variance can lead to more realistic ecological and evolutionary inferences ^135–138^. As with the trait data, we performed statistical validation analyses. We did this to identify the congruence between our data and that of the largest previously-assembled dataset of cactus distributions ^10^ (details in Technical Validation).

### Data Records

All data files described below are available in the GitHub repository (https://github.com/jamie-thompson/CactEcoDB) and Figshare (https://doi.org/10.6084/m9.figshare.30940019) ^139^. The dataset comprises cleaned species-level trait, spatial, environmental, phylogenetic, and diversification rate data for up to 1,063 species of Cactaceae (Figures 1, 2 and 5). Each file is formatted for ease of integration into ecological and evolutionary analyses.

1. **CactEcoDB_v1.csv**

Format: Spreadsheet (.csv) Description:

This file contains species-level data for Cactaceae, integrating trait, phylogenetic, spatial, and environmental information. Specifically, it includes:

● Mean tip-specific speciation rates estimated using BAMM, MiSSE and DR (lambda)
● Morphological and ecological trait data, such as maximum plant height (cm), growth form (binary and refined), pollination syndrome, and geographic range size (km^2^)
● Summary statistics (typically medians) of environmental variables based on georeferenced occurrence data

Fields:

● Diversification

○ BAMM_lamba_V1: Mean species-specific speciation rate estimated using BAMM ^97^ for phylogeny V1
○ BAMM_lamba_V2: Mean species-specific speciation rate estimated using BAMM for phylogeny V1
○ MiSSE_lamba_V1_full: Speciation rate estimated using MiSSE ^98^ for phylogeny V1
○ MiSSE_lamba_V1_pruned: Speciation rate estimated using MiSSE for phylogeny V1
○ MiSSE_lamba_V2_full: Speciation rate estimated using MiSSE for phylogeny V2
○ MiSSE_lamba_V2_pruned: Speciation rate estimated using MiSSE for phylogeny V2
○ DR_lambda_V1: Speciation rate estimated using the DR statistic ^35^ for phylogeny V1
○ DR_lambda_V2: Speciation rate estimated using the DR statistic for phylogeny V2
● Morphological and ecological traits

○ growth_form_binary: Coarse growth form classification (arborescent vs. compact group), following ^10,51^
○ growth_form_detailed: Detailed growth form description based on literature (arborescent, barrel, columnar, cushion forming, epiphytic, geophytic, globose caespitose, globose solitary, shrubby)
○ max_size_height_or_length: Maximum reported plant height or stem length (cm), building on ^10,51^
○ pollinator_binary: Pollination syndrome grouped into ancestral (bee) vs. derived (bat, bird, moth, other insects), according to ^10,51^
○ pollinator_detailed: Detailed pollination syndromes (bat, bird, moth, other insects)
○ chromosome_count_median: Median chromosome count (n)
○ chromosome_count_distribution: Summary information or distribution notes on chromosome counts (n) sourced from the Chromosome Counts Database ^123^
○ range_size: Geographic range size estimates (km^2^)
● Environmental variables (CHELSA bioclimatic variables ^132^, SoilGrids ^134^, and Global Multi-resolution Terrain Elevation Data (GMTED) ^133,140^)

○ Bio1–bio19: Standard bioclimatic variables, including temperature (bio1-bio11) and precipitation metrics, extracted from CHELSA ^132^. Bio1 to bio11 are temperature-related variables, and bio12-bio19 are precipitation-related. For example:

▪ bio1: Annual mean temperature (°C)
▪ bio2: Mean diurnal temperature range (°C)
▪ bio3: Isothermality (compares diurnal and annual temperature) (100 × (bio02 ÷ bio07))
▪ bio4: Temperature seasonality, the standard deviation of mean monthly temperatures (100/°C)
▪ bio5: Maximum temperature of warmest month (°C)
▪ bio6: Minimum temperature of warmest month (°C)
▪ bio7: Temperature annual range (°C)
▪ bio8: Mean temperature of wettest quarter (°C)
▪ bio9: Mean temperature of driest quarter (°C)
▪ bio10: Mean temperature of warmest quarter (°C)
▪ bio11: Mean temperature of coldest quarter (°C)
▪ bio12: Annual precipitation (kg m^-2^ year^-1^)
▪ bio13: Precipitation of wettest month (kg m ² month ¹)
▪ bio14: Precipitation of driest month (kg m ² month ¹)
▪ bio15: Precipitation seasonality (unitless): Coefficient of variation (100 × SD ÷ mean) of monthly precipitation
▪ bio16: Precipitation of wettest quarter (kg m ² month ¹)
▪ bio17: Precipitation of driest quarter (kg m ² month ¹)
▪ bio18: Precipitation of warmest quarter (kg m ² month ¹)
▪ bio19: Precipitation of coldest quarter (kg m ² month ¹)
▪ (See CHELSA documentation for full descriptions, especially with regards to scales and offsets used in some variables ^132^)
○ Aridity and productivity

▪ aridity_index: Aridity index (mean annual precipitation/potential evapotranspiration)
▪ pet: Potential evapotranspiration (kg m^-2^ year^-1^)
▪ npp: Net primary productivity (g C m^−2^ yr^−1^)
○ Elevation and topography extracted from GMTED ^133,140^

▪ elevation: Median elevation across occurrence records (m)
▪ slope: Median slope of the terrain (°)
▪ sd_slope: Median standard deviation (sd) of slope
▪ sd_elev: Median standard deviation of elevation
▪ roughness: Median terrain roughness (largest inter-cell difference of a focal cell and its 8 surrounding cells, m)
▪ prof_curve: Median standard deviation of profile curvature (sd of radians per meter)
▪ tang_curve: Median standard deviation of tangential curvature (sd of radians per meter)
○ Soil Variables extracted from SoilGrids ^134^

▪ soil_sand: Median proportion of sand particles (> 0.05/0.063 mm) in the fine earth fraction
▪ soil_water: Median derived available soil water capacity (volumetric fraction)
▪ texture: Soil texture class, classified into twelve categories (e.g. sand, silt, clay, loam, silty clay)

Notes:

● Speciation rates estimated with BAMM represent mean species-specific estimates from the entire posterior sample. The full BAMM posterior event data objects are available separately in CactEcoDB, and can be analyzed (e.g. rates through time and deep rate shifts) using the R package BAMMtools ^106^.
● Plant height values are not statistical species means from population samples, but are derived from maximum reported sizes in taxonomic or ecological literature, as is common across plant research generally ^10,110,141–144^.
● Pollinator data was rarely specific to individual species of pollinator. Therefore, we classified them into major taxonomic groups described in ^10,51^ as is standard practice in pollinator research ^51,111–114^.

2. **CactEcoDB_coords_envs.csv**

Format: csv

Description: Curated distribution coordinates with associated environmental values for every coordinate.

Fields: species, decimal latitude, decimal longitude, environmental variables (e.g. bio1-19, aridity index, soil texture).

Notes: Most of these coordinates are randomly sampled from expert IUCN ranges, some of them are curated GBIF occurrences. We have written permission from the IUCN Red List to reproduce randomly-sampled coordinates from expert ranges.

3. **CactEcoDB_phylogeny_V1.tre**

Format: Newick (.tre)

Description: Time-calibrated phylogeny for 1,063 Cactaceae species, inferred using a supermatrix approach and Penalized Likelihood dating.

Notes: Outgroups are pruned and tips are matched to species names in the trait and spatial datasets to facilitate research.

4. **CactEcoDB_phylogeny_V2.tre**

Format: Newick (.tre)

Description: Time-calibrated phylogeny for 1,052 Cactaceae species, inferred using a supermatrix approach and Penalized Likelihood dating.

Notes: This differs from the V1 Phylogeny, in that is uses a phylogenomic backbone to improve relationships ^89^.

5. **CactEcoDB_concatenated_alignments.fasta**

Format: Fasta (.fasta)

Description: concatenated alignment of 18 loci, first used in Thompson et al. ^10^.

Notes: Locus regions are defined in CactEcoDB_phylo_partition.txt.

6. **CactEcoDB_nucleotide_accessions.csv**

Format: csv

Description: A species-by-locus matrix of GenBank accessions used to make both phylogenies. Originally assembled by Thompson et al. ^10^

7. **CactEcoDB_phylo_partition.txt**

Format: RAxML-format partition file (.txt) Description: Defines the start and end of loci in

CactEcoDB_phylo_concatenated_alignment.fasta, used to apply a different nucleotide substitution model to each locus.

8. **CactEcoDB_phylogeny_V1_bootstrap.tre**

Format: Newick (multiple phylogenies)

Description: The sample of 300 bootstrap replicates estimating alongside the maximum likelihood search for phylogeny V1 estimated by Thompson et al. ^10^

Notes: This can be used to assess topological support and replicate analyses while accounting for phylogenetic uncertainty

9. **CactEcoDB_phylogeny_V2_bootstrap.tre**

Format: Newick (multiple phylogenies)

Description: The sample of 300 bootstrap replicates estimating alongside the maximum likelihood search for phylogeny V2

Notes: This can be used to assess topological support and replicate analyses while accounting for phylogenetic uncertainty

10. **CactEcoDB_BAMM_eventdata_V1.txt**

Format: txt

Description: The posterior sample of rate shift configurations (eventdata) estimated by BAMM.

Fields: specific to BAMM ^97^, use the R package BAMMtools ^106^ to access and analyse this.

Note: We do not provide this for MiSSE and DR, for which we provide tip-speciation rates in the main CactEcoDB file. BAMM is a flexible system for rates through time and rate shifts, therefore we provide the intermediary event data file. Tip-speciation rates calculated from BAMM event data files are already present in the main CactEcoDB file.

11. **CactEcoDB_BAMM_eventdata_V2.txt**

Format: txt

Description: The posterior sample of rate shift configurations (eventdata) estimated by BAMM.

Fields: specific to BAMM ^97^, use the R package BAMMtools ^106^ to access and analyse this.

12. **CactEcoDB_variable_descriptions.txt**

Format: txt

Description: Definitions, units, and data sources for all variables in the dataset. Notes: Intended to support reproducibility and proper variable interpretation.

13. **CactEcoDB_full_sources_shorthand.xlsx**

Format: xlsx

Description: References (in shorthand, see CactEcoDB_full_references_lookup.xlsx file) for all variables in CactEcoDB.

Notes: Intended to support transparency, traceability and reliability.

14. **CactEcoDB_full_references_lookup.xlsx**

Format: xlsx

Description: A lookup table matching the shorthand reference style (e.g. IUCN, CHELSA, BAMM) with full references.

Notes: Intended to support transparency, traceability and reliability.

15. **CactEcoDB_code.zip**

Format: compressed folder (ZIP) of code scripts

Description: Contains scripts (primarily R) for data assembly, random sampling of spatial coordinates, taxonomic and spatial distribution cleaning, estimation of diversification rates, post-processing (e.g. for BAMM), outlier detection, and visualisation.

### Technical validation

To ensure the robustness and reliability of CactEcoDB, we implemented multiple validation steps integrating automated and manual approaches, which we applied to the phylogenies, diversification estimates, trait data, species distributions, and environmental summaries.

### Phylogenies and diversification rate estimates

It is well known that phylogenetic relationships in cacti are difficult to resolve due to their low sequence divergence ^10,51,52,55,60,89^. There are different approaches available, such as constructing phylogenies for individual groups (e.g. genera and tribes ^64,65,145^), deeply sampling entire genomes to maximise backbone resolution ^89,146^, and the supermatrix approach, which maximises species sampling at the cost of the support provided by genome-wide data ^21,147–150^. Here, we have provided preliminary phylogenies using the supermatrix approach, in order to maximise species sampling for comparative research, but note that caution should be taken, as with all supermatrix phylogenies ^151–155^. The phylogenies were constructed using 18 multiple sequence alignments of commonly sequenced nucleotide loci publicly available in Genbank. All alignments were manually and statistically inspected, and corrected for quality and taxonomic coverage (see ^10^).

Maximum Likelihood phylogenetic reconstruction was performed with partitioned models to account for variation in substitution rates between loci, and bootstrap support was calculated. In V1, a relaxed constraint was used to improve the likelihood computation and in V2, we made use of a recently-published phylogenomic tree of cacti ^89^, in which we implemented a much stricter topological constraint. Using both a largely data-driven reconstruction (V1) and a topology guided by a recent phylogenomic framework (V2) provides complementary strengths. V1 maximises the signal present in the sequence data, while V2 enforces well-supported relationships among deeper nodes, thereby improving overall phylogenetic stability. For users of the data wishing to explore the phylogenetic uncertainty, we provide 300 bootstrap replicates for each topology, as well as the full alignment and partition file. Time-calibration used secondary divergence estimates, as there is no useful fossil record for cacti ^55^. Resulting time-calibrated phylogenies are the widest-sampled for cacti, encompassing all major lineages, although sampling remains sparse in some poorly-sequenced lineages. Despite incomplete sampling and inherent limitations of time-calibration, the topologies and divergence timings are broadly consistent with other recent reconstructions, which increasingly converge on a crown age of ∼27-37 Mya for the family ^5,51,55,146^. Our phylogenies, like all divergence-time estimates lacking direct fossil calibration, carry uncertainty in absolute node ages; however, their relative branch lengths and overall temporal structure are robust and well suited for comparative analyses such as tip-rate estimation, diversification modelling, and phylogenetic correction ^10^.

We provided multiple estimates of diversification rate to account for methodological and topological differences. We provided BAMM estimates, followed established statistical protocols when setting conservative priors ^106^, accounted for imbalance sampling efforts at a high resolution of genus-level (fractions available at ^10^), and ensured convergence was reached. We provided MiSSE estimates as an alternative ^98^, which allows users to test whether results change according to method, especially differences in the implementation of corrections for incomplete taxonomic sampling. We also provide the DR statistic ^35^, which makes fewer assumptions regarding the evolutionary model underlying phylogenetic branching, but is noisy and cannot account for incomplete sampling. Diversification rates estimated with MiSSE used a global sampling fraction to account for incomplete sampling, and were repeated with and without the shortest branches. We tested the congruence of all eight estimates (four per phylogeny: BAMM, DR statistic, MiSSE with all species and MiSSE without short terminal branches).

Pairwise Pearson correlations of log-transformed tip rates revealed strong and consistent agreement across approaches. All pairwise comparisons were strongly significant (p < 0.000001), and correlation coefficients ranged from 0.24 to 87 (mean r = 0.61, median r = 0.64). We advise that researchers use multiple approaches to investigating diversification rates, with an understanding of the uncertainties and methodological differences between methods.

### Trait data

As with similar trait datasets ^14,25,26,28,29,156^, when collecting trait data, we relied on authoritative textbooks and peer reviewed literature, using secondary sources (e.g. online floras) infrequently, and only when it provided a sensible value (i.e. not when photographs of a cactus is clearly miniature but the recorded plant height is unrealistically large). We used standardised measurement protocols to minimise errors, such as unit conversion errors that are present in some databases.

Once collected, trait data were checked for plausibility and consistency through a combination of automated flagging, literature cross-checking and expert review. Generally, minimum and maximum plant height values were most frequently recorded and individual level variation is not available, as is common in other groups ^17,42,141^. We retained maximum values, as is best-practice in comparative research on plant height to avoid issues associated with diseased, juvenile or incomplete specimens ^110,141^. Height values that were extreme (e.g. 0cm, 2500cm) or contradictory were flagged and verified against alternative sources when available. Growth form data were collected and discussed between authors in accordance with our defined groupings. We flagged records with outlying plant height for their growth form, on the principle that growth form shapes height (i.e. globose species are much smaller than arborescent species, and shrubby species and somewhere between). Outlier detection was performed within each growth form, and polymorphic species were expanded so that their height values were evaluated against every growth form they express. We defined outliers following the standard interquartile method and inspected 75 that were detected. Of these, 11 were removed, three were changed, and 61 were kept as-is. Following this, an author with 30 years experience in growing and over 30 field work trips to diverse cactus ecosystems (AG), checked all the data. Pollinator data required significant additional validation. A portion in Thompson et al. ^10^ originated from Hernández-Hernández et al. ^51^, where many pollination syndromes were provided without traceable primary sources. To assess the reliability of these data, we searched independently for evidence for each species using published ecological studies, species-level monographs, and an alternative collection which recorded sources ^120^, the latter of which we incorporated into CactEcoDB. Where independent sources were available, we updated or confirmed the pollinator data accordingly. For the 56 cases where no primary or secondary documentation could be found, we removed these from the dataset. The retained categories follow the definitions widely used in previous macroevolutionary work on Cactaceae ^51,60^.

As a final validation step for plausibility, one author (AG), who has over 30 years experience in the cactus horticulture industry and has undertaken 30 field trips to visit diverse cactus habitats in the Americas, checked all trait data one by one. This expert review step identified additional implausible datapoints that were not feasible by statistical testing and literature checkers. Notably 28 species were flagged by AG, many of which because their heights were likely misrecorded by Anderson ^109^ based solely on single stem segment measurements. A similar issue was identified in Anderson ^109^ for one species (*Opuntia chaffeyi*), where the reported height is ambiguously recorded and may correspond to the size of a rhizome instead of the entire plant. These were not possible to verify independently and were removed.

As with all comparative datasets, especially for plants ^13,42,47,157^, there are gaps owing to imbalanced data collection efforts. The poorest-covered traits are pollinator and chromosome count, because these are difficult to collect in the field compared to growth form and plant height descriptions. Although we note that for pollinators, CactEcoDB provides observations for a large proportion of species when compared to other recent datasets for plant families (17.68% of species compared to ∼10% for Orchidaceae ^114^). Gaps in pollinator observation data are fairly group-specific; there are relatively more species with data in Cactoideae than in Opuntioideae, for example, and more in Cactoideae tribes Cereeae and Phyllocacteae, than Cacteae. In contrast, chromosome count data collection efforts seem less biased by group, and both growth form and plant height are fairly well covered as commonly-recorded variables.

### Spatial distributions and environmental variables

Spatial distributions were made of a combination of high resolution expert-ranges (iucnredlist.org) and a smaller number of curated occurrences where expert ranges were unavailable, following established protocols ^128^. Randomly sampled coordinates from expert ranges were visually inspected per species, which identified one unlikely range that is the result of an introduced population. Occurrences from GBIF were thoroughly checked following established protocols ^109,129,130^, which performed a six step procedure, flagging coordinates based on spatial and statistical issues ^131^, and then filtering according to independent estimates of distributions, broadly corresponding to country scale ^88^. These two complementary approaches maximise coverage and reliability. Expert-defined range polygons provide consistent distributions for most species, while rigorously filtered occurrence data extend sampling to species lacking expert maps. As with all large-scale distribution datasets, ranges should be viewed as the best current synthesis and may change as taxonomy is revised, population shifts are documented, introductions are identified, and ongoing IUCN assessments and occurrence data updates redefine the distributions. As with the phylogenies, we will update the distributions in the future as new data compilations emerge.

Environmental variables were extracted from high-resolution raster datasets (e.g. CHELSA ^132^, SoilGrids ^134^ for each coordinate, and species-level summaries were calculated as medians to minimise the influence of outliers following ^10^. Critically, we statistically compared values of extracted spatial data between previous methods that employed GBIF-derived data ^10^ and our current method, and found very strong agreement, demonstrating that our approach of sampling from expert IUCN ranges is appropriate. We estimated 1. linear regressions and 2. Pearson correlation coefficients between variables in CactEcoDB and in the largest previous spatial dataset for cacti (Thompson et al., 2024). We found that comparisons between all 31 environmental variables are significant at p < 0.05, with a median slope and correlation coefficient of 0.89 and 0.88, respectively. Furthermore, 25 comparisons recovered a slope and correlation coefficient of > 0.75. Our new approach also retrieved valid data for soil texture, which is discrete and has strong agreement (72.60%) with data retrieved from GBIF occurrences (Thompson et al., 2024).

### Usage Notes

CactEcoDB is the first large-scale integrative database for the cactus family, and brings together multiple data types that are rarely available in combination for any plant family, including species-level traits, expert-curated spatial distributions, environmental variables, time-calibrated phylogenies and speciation rate estimates. Such a synthesis has not been previously available for cacti, despite their interest to biologists due to their exceptional morphological and ecological diversity, and status as one of the most threatened groups worldwide ^70^. By assembling these diverse data sources, CactEcoDB enables comparative, macroecological, macroevolutionary and conservation analyses that were previously impractical or impossible due to fragmented data.

As is typical of large-scale biodiversity datasets, CactEcoDB reflects uneven data availability across species and data types ^47,158^. Trait coverage is extensive across the family, but completeness varies by trait and specific group. For plant height, values were derived from sources that only reported minimum and maximum height. In these cases, maximum values were retained to reduce bias from juvenile, incomplete, or diseased specimens. However, these records often lack associated metadata (e.g., number of individuals measured, habitat context), and users should treat them as approximations of mature phenotypes rather than statistically derived species means. The same can be true for chromosome count, which can vary intraspecifically.

CactEcoDB is designed to be a continually evolving resource. As new trait measurements, spatial distributions, molecular sequences, and relevant environmental maps become available, we will incorporate these into future versions. We therefore welcome contributions of additional data and corrections to further improve the accuracy, completeness, and utility of CactEcoDB.

In summary, CactEcoDB offers a comprehensive resource for large-scale ecological and evolutionary analyses, but as with any aggregated dataset, its contents should be interpreted in light of the inherent variability in data completeness and precision. Where greater taxonomic or spatial resolution is required, users may find it helpful to consult original sources or species-level references. Similar to other trait databases ^15,42,157,159–162^, we recommend that users monitor ongoing changes in Cactaceae taxonomic nomenclature.

## Code availability

Code used to generate, process and analyse data, and produce figures is available at Figshare (https://doi.org/10.6084/m9.figshare.30940019) ^139^ and in the GitHub repository (https://github.com/jamie-thompson/CactEcoDB).

## Acknowledgments

We acknowledge Craig Hilton-Taylor for granting permission to reproduce IUCN spatial coordinates, Eric Hagen for technical advice running MiSSE, Juriaan de Vos for advice on the phylogenomic backbone, and Itay Mayrose for granting permission to reproduce chromosome counts from the Chromosome Count Database.

